# VirtualSkin integrates skin-cell identity with intervention-response prediction

**DOI:** 10.64898/2026.10.06.756877

**Authors:** Mira Velland, Elias Norwick, Sofia Ardene, Theo Marwick, Lina Corven, Adrian Solmere, Nia Calder, Rafael Quillon, Isobel Renn, Marcus Elian

**Author notes:** Correspondence: Marcus Elian. These authors contributed equally (Mira Velland, Elias Norwick).

## Abstract

Human skin atlases resolve a diverse set of cellular states, yet connecting these states to their responses to intervention remains difficult across independently sampled tissues. Here we present VirtualSkin, a cell–language model that aligns skin transcriptomes with biological descriptions and supports intervention-conditioned response prediction. We evaluated the model across independent atlas, disease, anatomical and perturbation cohorts, with donors, studies and perturbations withheld according to the experimental design. VirtualSkin improved inflammatory-skin annotation by 0.056 macro-F1 points and cell–text retrieval by 0.12 Recall@10 points relative to the strongest corresponding comparators. The representation retained cell identity during study integration and transferred across ageing, systemic sclerosis, lymphoma and anatomical sites. Across five intervention cohorts, response prediction improved by 0.03–0.09 response-score points, with agreement at the levels of gene-expression change, gene recovery and transcriptional programs. Paired tissue trajectories provided a complementary assessment in two donors per intervention. Component ablations and pairing controls localized the predictive contributions, while uncertainty estimates supported selective prediction under distribution shift. These results connect skin-cell recognition to intervention-response prediction within a common evaluation framework and define the conditions under which predictions can inform experimental prioritization.

## Introduction

Human skin combines a renewing epithelial barrier with stromal, immune and vascular compartments that respond together to injury and disease. The identities of these cells depend on developmental stage, anatomical site and their local tissue environment. Single-cell studies have resolved epidermal differentiation states and disease-associated fibroblast populations, while spatial profiling has placed these states within inflammatory and tumour niches.^1,2,3,4,5,6,7^ This accumulating detail creates an opportunity to model cutaneous biology at cellular resolution. It also exposes a gap between describing a cell’s present state and anticipating how that state changes after intervention.

The gap is particularly consequential in skin. A fibroblast program associated with fibrosis may resemble a program found in repair, yet the response to a defined treatment can depend on the donor, tissue site and initiating mechanism. Likewise, tumour-associated and inflammatory immune states can share markers while differing in their response to cytokine exposure, gene disruption or drug treatment.^8,9,10,11,12,13^ An informative representation must therefore preserve distinctions within a lineage and remain interpretable across tissue contexts. Accurate assignment of a broad cell label is an intermediate requirement; response prediction additionally requires a model to recover changes that were unavailable from the baseline state alone.

Single-cell foundation models offer reusable representations learned from large collections of transcriptomes.^14,15,16,17^ Cell–language approaches extend these representations by linking expression to descriptions, enabling annotation and retrieval with less direct supervision.^18,19^ Models that combine dissociated and spatial data further connect molecular identity to tissue context.^20^ In parallel, perturbation models predict transcriptional outcomes from genetic or chemical interventions, including combinations that are absent from training.^21,22,23^ The integration of these capabilities in skin requires a common set of comparisons that follows the progression from molecular representation to response prediction.

Several properties of public skin datasets make such comparisons demanding. Studies differ in cell capture, anatomical coverage, clinical composition and the resolution of their annotations. Multiple cells from the same donor provide detailed measurements without providing the same number of independent biological replicates. Text can carry a similar dependence: a model may retrieve an appropriate description because the query and target repeat the same label or originate from the same publication. Atlas benchmarks accordingly distinguish biological conservation from batch removal, and sample-aware models explicitly retain donor variation.^24,25,26,27,28,29^ Perturbation benchmarks impose an additional requirement that the response of the held-out intervention is unavailable during fitting and model selection.^23,30^

We developed VirtualSkin to connect cutaneous cell identity, biological language and intervention response. The evaluation asks three successive questions: whether the representation preserves skin-cell states across study boundaries; whether it supports annotation, retrieval and external transfer when supervision or input quality is limited; and whether the resulting model recovers measured responses to held-out interventions. We address these questions with cohort-resolved comparisons, matched method inputs, and controls for molecular, lexical and experimental shortcuts. Figure 1 connects cohort provenance to the representation architecture and training objectives; Figure 2 defines the independent evaluation framework; Figures 3–6 test recognition and transfer; Figures 7–9 examine response accuracy, molecular fidelity and reliability. This sequence links each broader claim to the comparison needed to establish it.

**Fig. 1.**
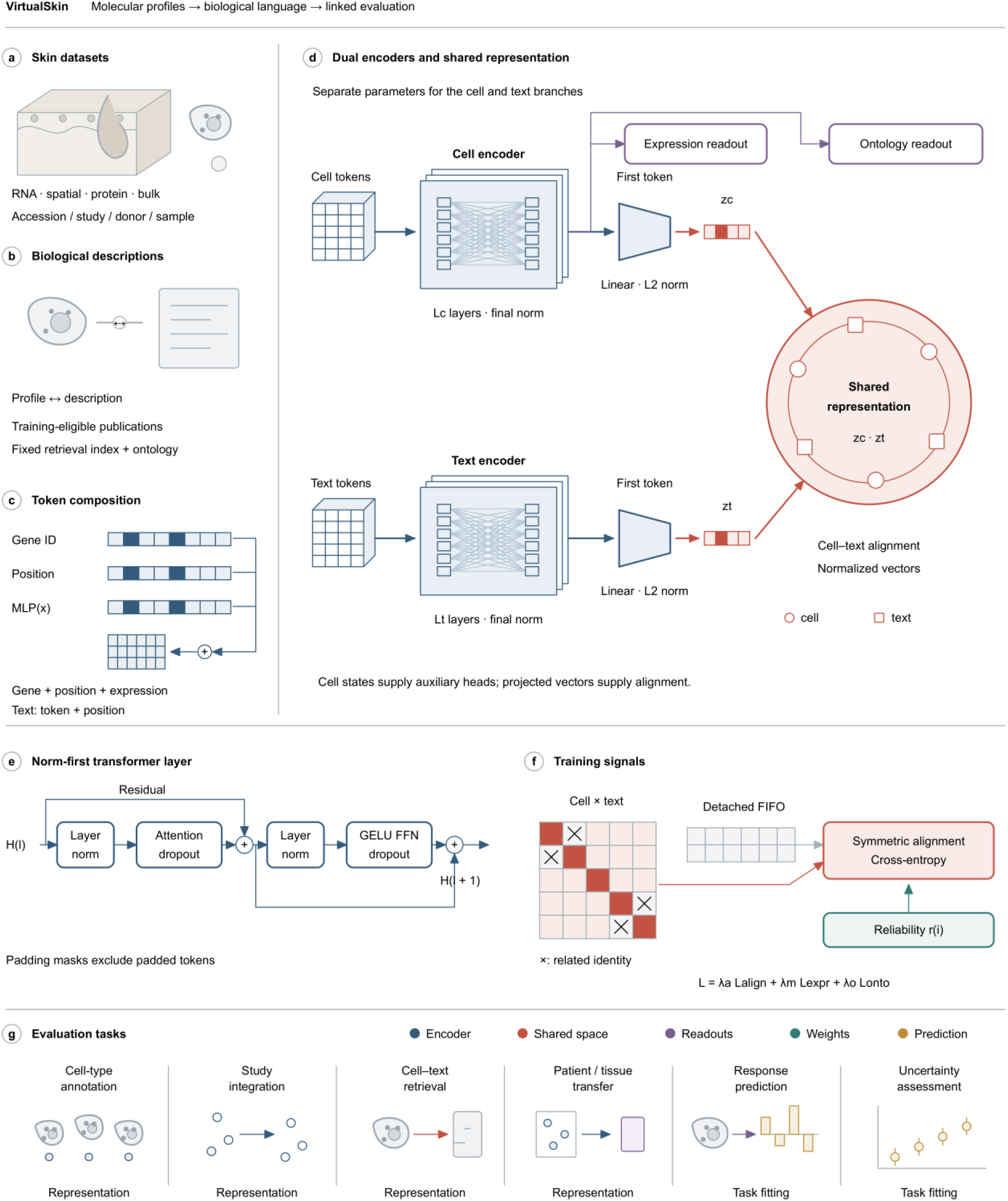
VirtualSkin inputs, representation architecture and training objectives. **a**, Accession-linked cutaneous studies are reconciled at study, donor and sample levels; RNA cells or nuclei, spatial sections and protein or bulk measurements retain separate assay denominators. **b**, Molecular profiles are paired with provenance-linked biological descriptions from training-eligible publications; related cell identities supply the declared ontology. **c**, Gene, position and projected scalar-expression embeddings are added to form cell tokens. Text inputs combine token and position embeddings. **d**, Separate cell and text transformers produce normalized, linearly projected first-token representations that converge in the shared representation. Circles and squares indicate cell and text vectors, respectively. The cell hidden states supply masked-expression reconstruction, and the pooled cell state supplies ontology classification. Lc and Lt denote encoder depths. **e**, A norm-first transformer layer uses multi-head attention, a GELU feed-forward block, dropout and residual additions; padding masks exclude padded tokens from attention. **f**, Symmetric cell-to-text and text-to-cell alignment uses pair-reliability weights. Related nonmatching labels are masked within the batch while matched pairs remain on the diagonal. Detached FIFO representations extend the negative set. The total objective combines alignment, smooth L1 expression reconstruction and ontology cross-entropy with the declared weights. **g**, The representation supports annotation, integration, retrieval and transfer; response and uncertainty are evaluated under their task-specific fitting specifications. Colour identifies the encoder, shared representation, auxiliary readouts, reliability weights and predicted readouts. Encoder widths, depths and training configuration are given in Methods and Supplementary Table 8.

**Fig. 2.**
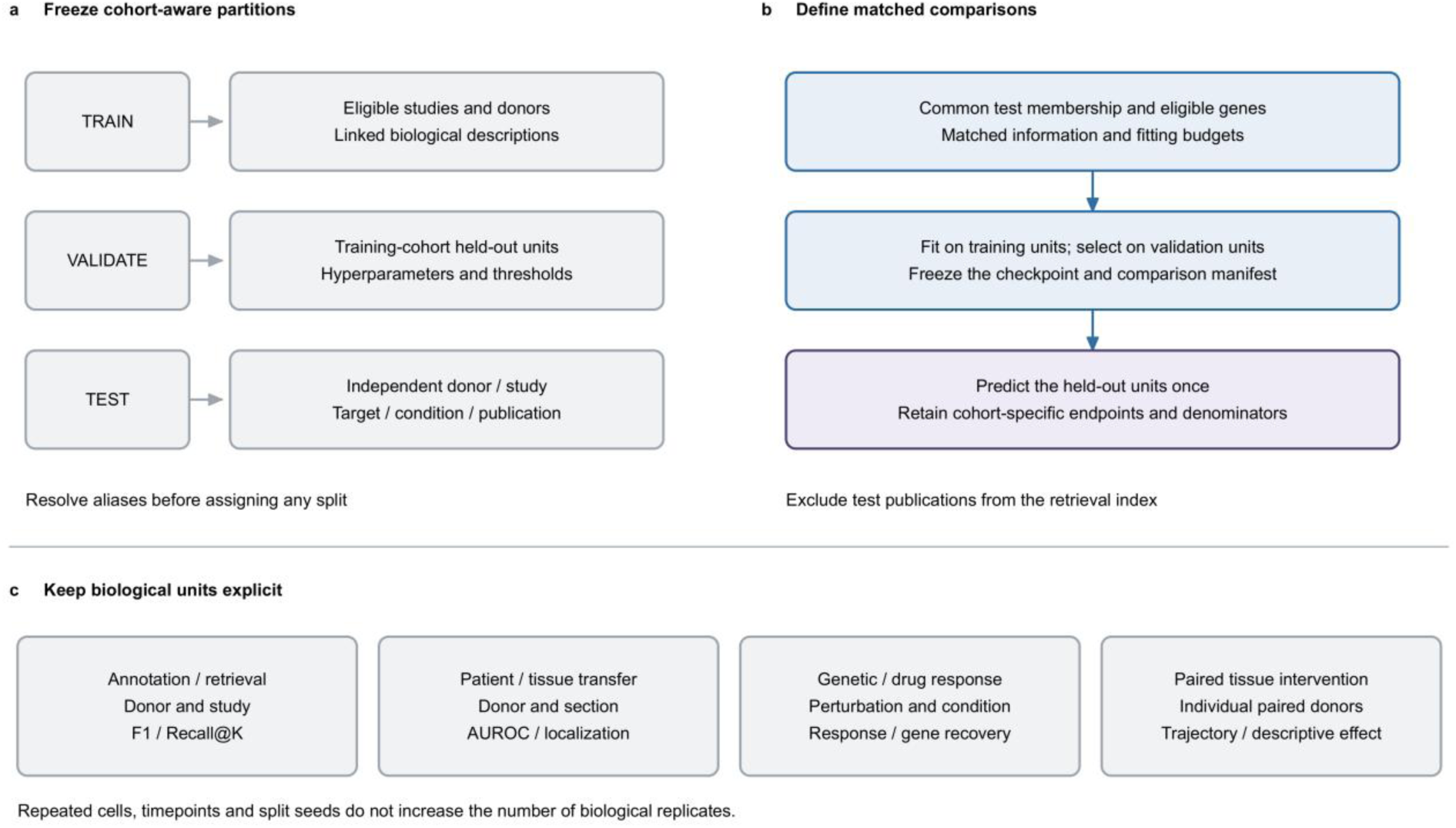
Cohort-resolved fitting and independent evaluation. **a**, Cross-portal aliases are resolved before assigning training, validation and test units. Held-out units follow each cohort’s donor, study, intervention or publication structure. **b**, Compatible methods share eligible test membership, genes and information. Model fitting uses training units; hyperparameter and threshold selection uses validation units. The checkpoint and comparison manifest are frozen before test prediction, and test publications are excluded from the retrieval index. **c**, Annotation and retrieval are evaluated across donors and studies, patient and tissue transfer across donors and sections, and genetic or drug response across perturbations or conditions. The two tissue-intervention cohorts retain individual paired-donor trajectories. Repeated cells, timepoints and split seeds do not increase the number of biological replicates. Endpoint definitions and eligible comparisons are provided in Supplementary Table 2.

**Fig. 3.**
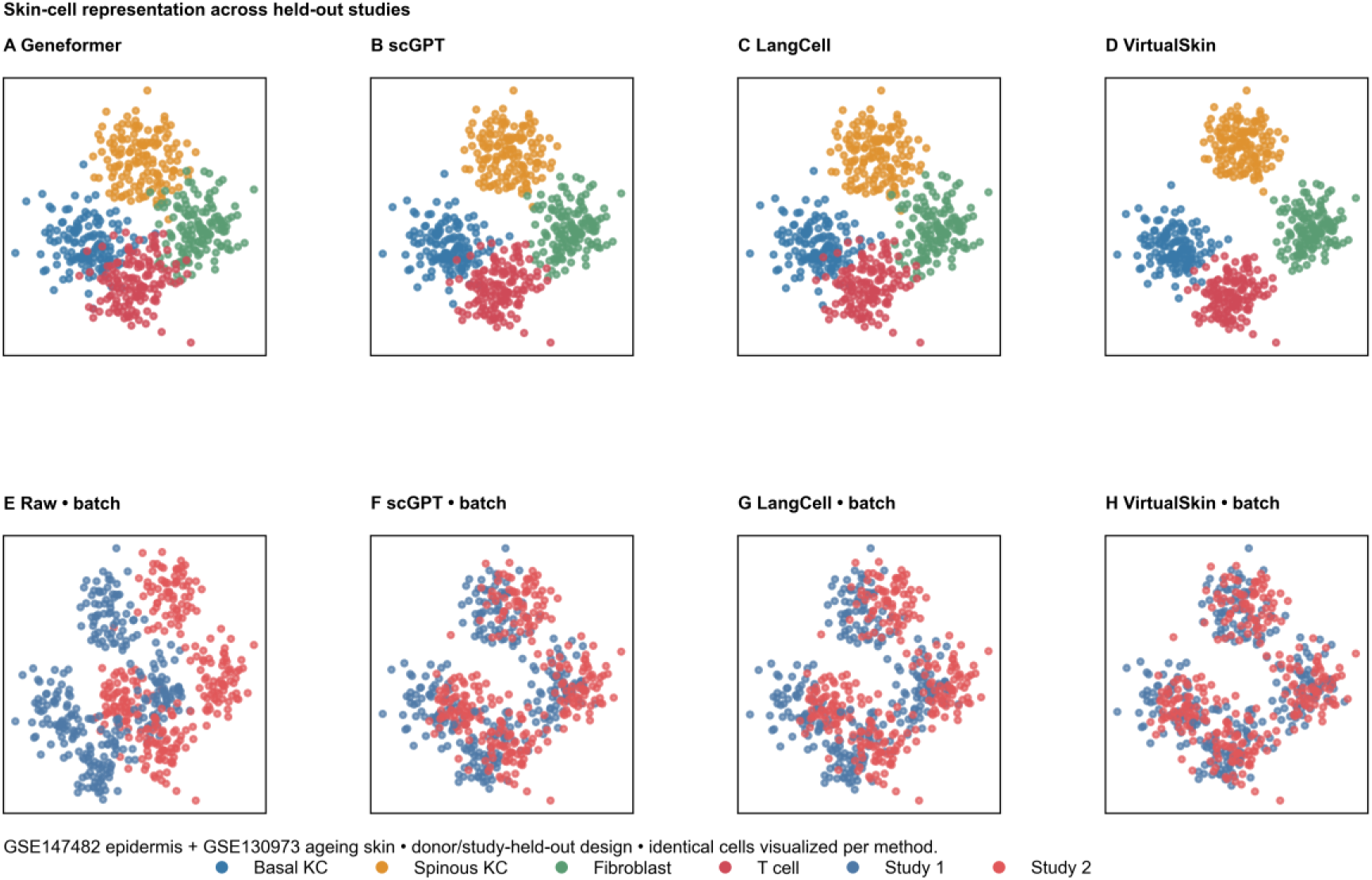
Skin-cell identity and study integration. **A–D**, Two-dimensional projections of the same displayed cells from GSE147482 and GSE130973 using Geneformer, scGPT, LangCell and VirtualSkin, respectively. Colours denote basal keratinocytes, spinous keratinocytes, fibroblasts and T cells. **E–H**, Study-coloured displays before transformation and after scGPT, LangCell and VirtualSkin transformation, respectively. The panels provide two-dimensional views of cell-class and study structure; annotation and transfer endpoints are evaluated in subsequent figures.

**Fig. 4.**
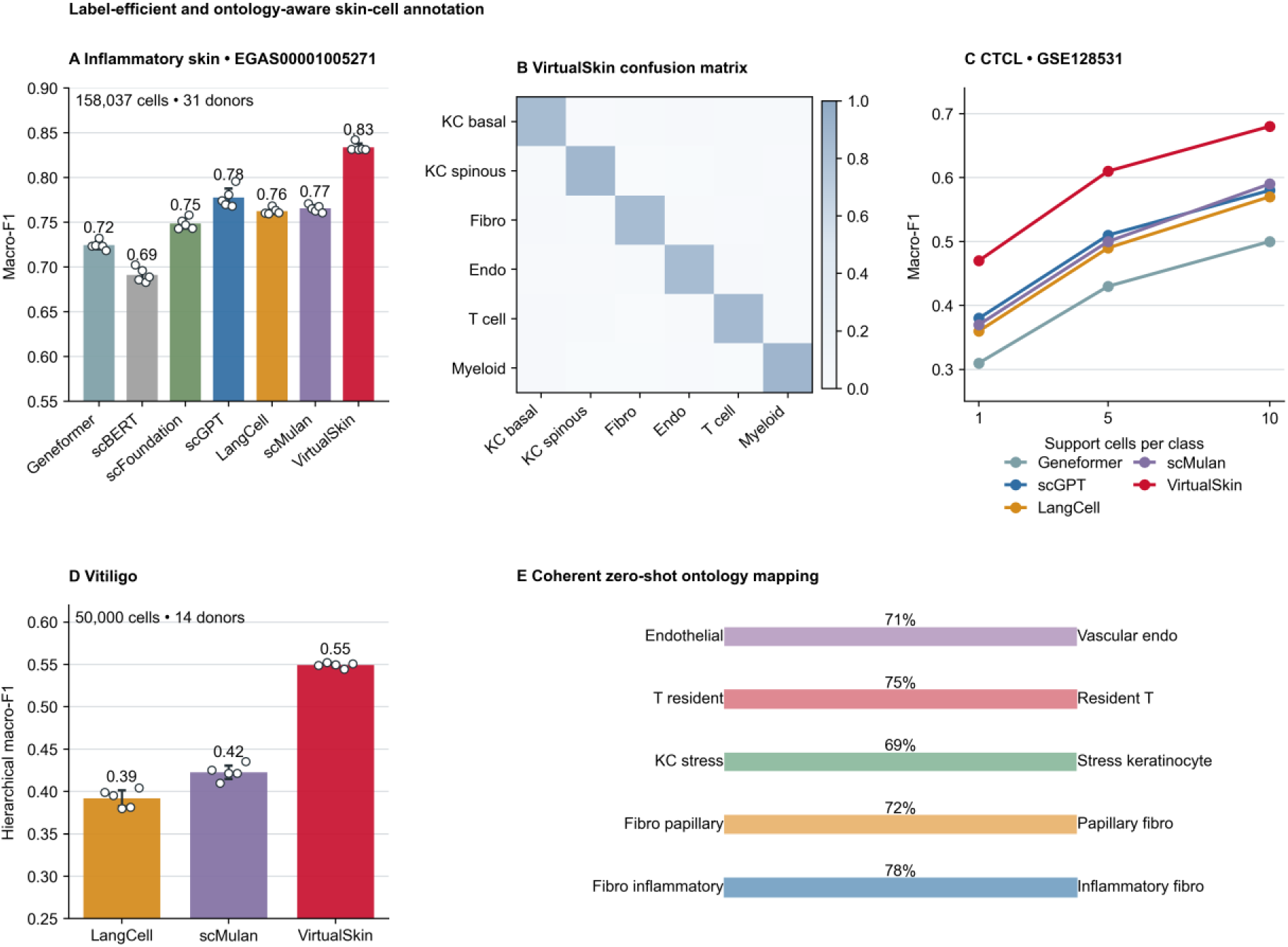
Annotation with limited supervision and transferred labels. **A**, Inflammatory-skin macro-F1 in EGAS00001005271 for seven representation methods. Bars denote the mean across five fixed splits; points denote individual split estimates. **B**, Row-normalized VirtualSkin confusion matrix for six broad cell groups; rows indicate reference labels and columns predicted labels. **C**, Cutaneous T-cell lymphoma macro-F1 in GSE128531 with one, five or ten labelled support cells per class. **D**, Hierarchical macro-F1 in vitiligo for LangCell, scMulan and VirtualSkin. **E**, Five source–target ontology correspondences; percentages denote the fraction of eligible source-label predictions assigned to the displayed target term. Error bars in **A,D** show the mean plus or minus 1.96 standard errors across the five displayed estimates. Class, donor and split counts are distinguished in Supplementary Table 4.

**Fig. 5.**
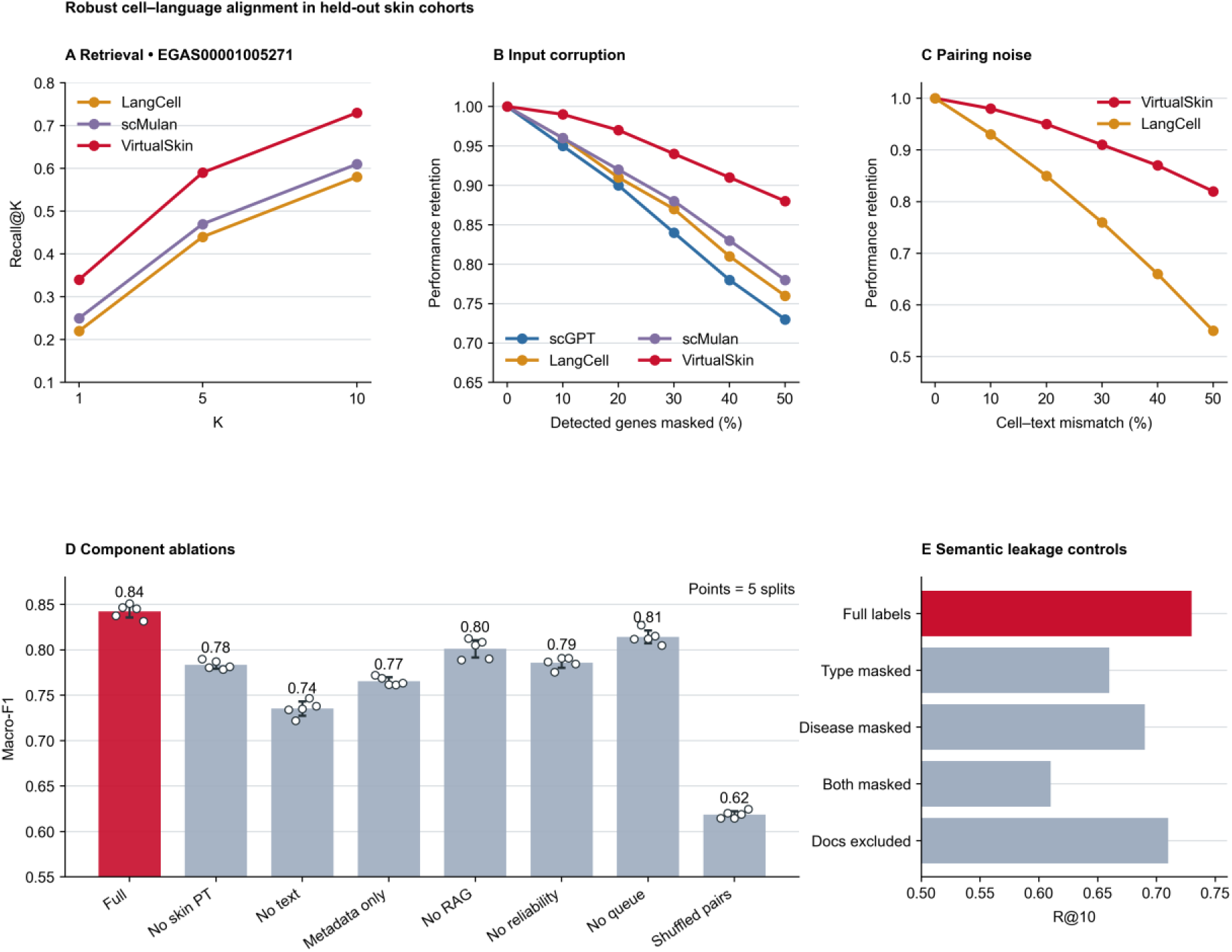
Cell–language retrieval, corruption and component controls. **A**, Cell-to-text Recall@1, Recall@5 and Recall@10 in held-out inflammatory skin for LangCell, scMulan and VirtualSkin. **B**, Performance retained after masking 0–50% of detected genes, relative to each method’s unmasked score; each curve is normalized to its own unmasked endpoint. **C**, Retention after replacing 0–50% of cell–description pairs. **D**, Annotation macro-F1 for the full model and seven component or pairing configurations. Bars show the mean across five splits, points individual split estimates and error bars the mean plus or minus 1.96 standard errors across those estimates. **E**, Recall@10 with full descriptions, cell-type terms masked, disease terms masked, both masked or benchmark-publication documents excluded. Candidate pools and corruption assignments are shared across methods within each comparison. RAG denotes retrieval augmentation.

**Fig. 6.**
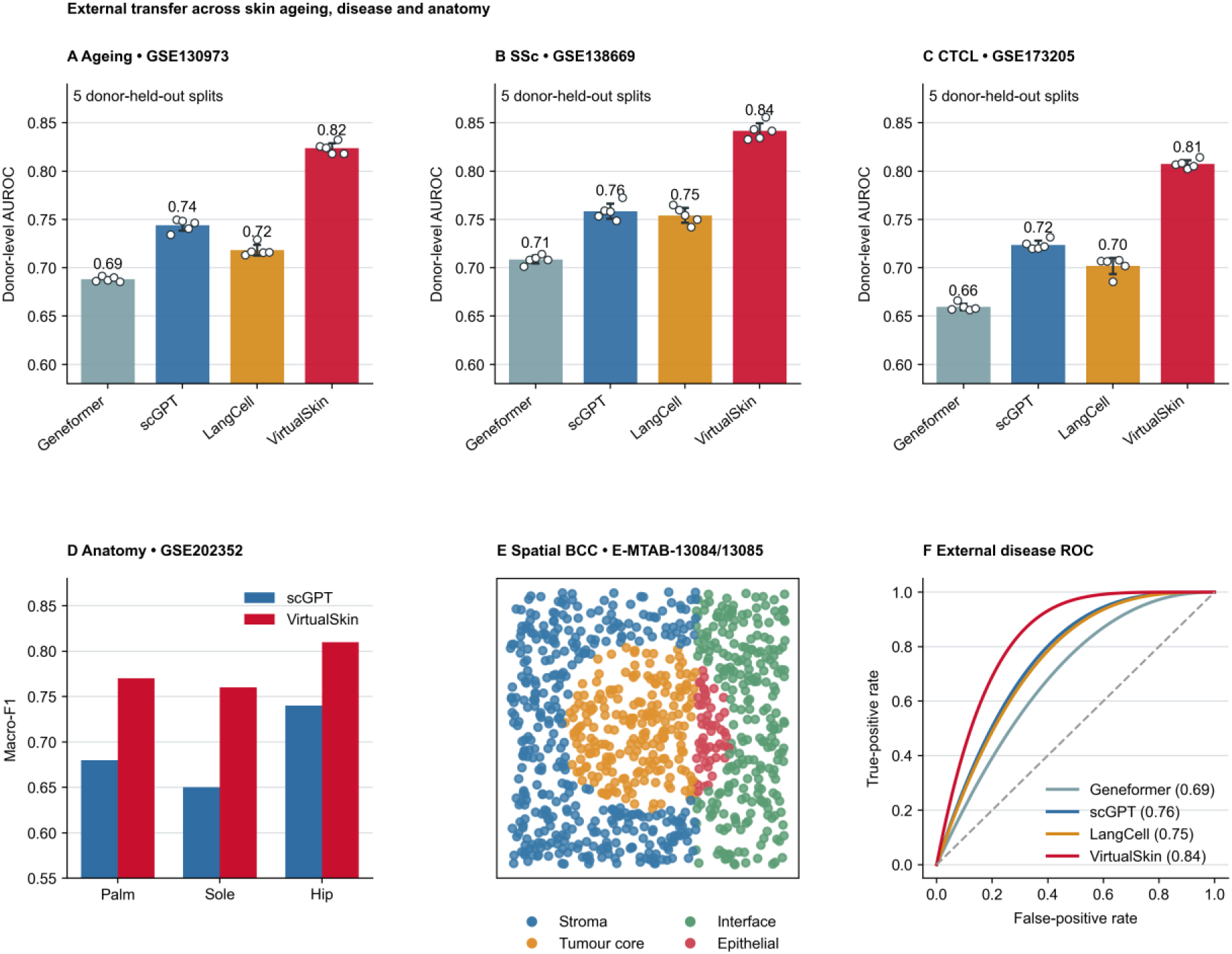
External age, disease, anatomical and spatial transfer. **A–C**, Donor-level AUROC in ageing skin (GSE130973), systemic sclerosis (GSE138669) and cutaneous T-cell lymphoma (GSE173205), respectively. Bars show the mean across five fixed splits, points individual split estimates and error bars the mean plus or minus 1.96 standard errors across those estimates; cohort endpoint definitions are in Supplementary Table 2. **D**, Annotation macro-F1 in palm, sole and hip skin from GSE202352 for scGPT and VirtualSkin. **E**, Tissue compartment assignments in basal-cell carcinoma spatial assays E-MTAB-13084 and E-MTAB-13085, coloured as stroma, tumour core, interface or epithelium. **F**, ROC curves from the same donor-level score construction for the designated external disease endpoint. The endpoint, operating rule and donor counts are reported in Supplementary Tables 2 and 3.

**Fig. 7.**
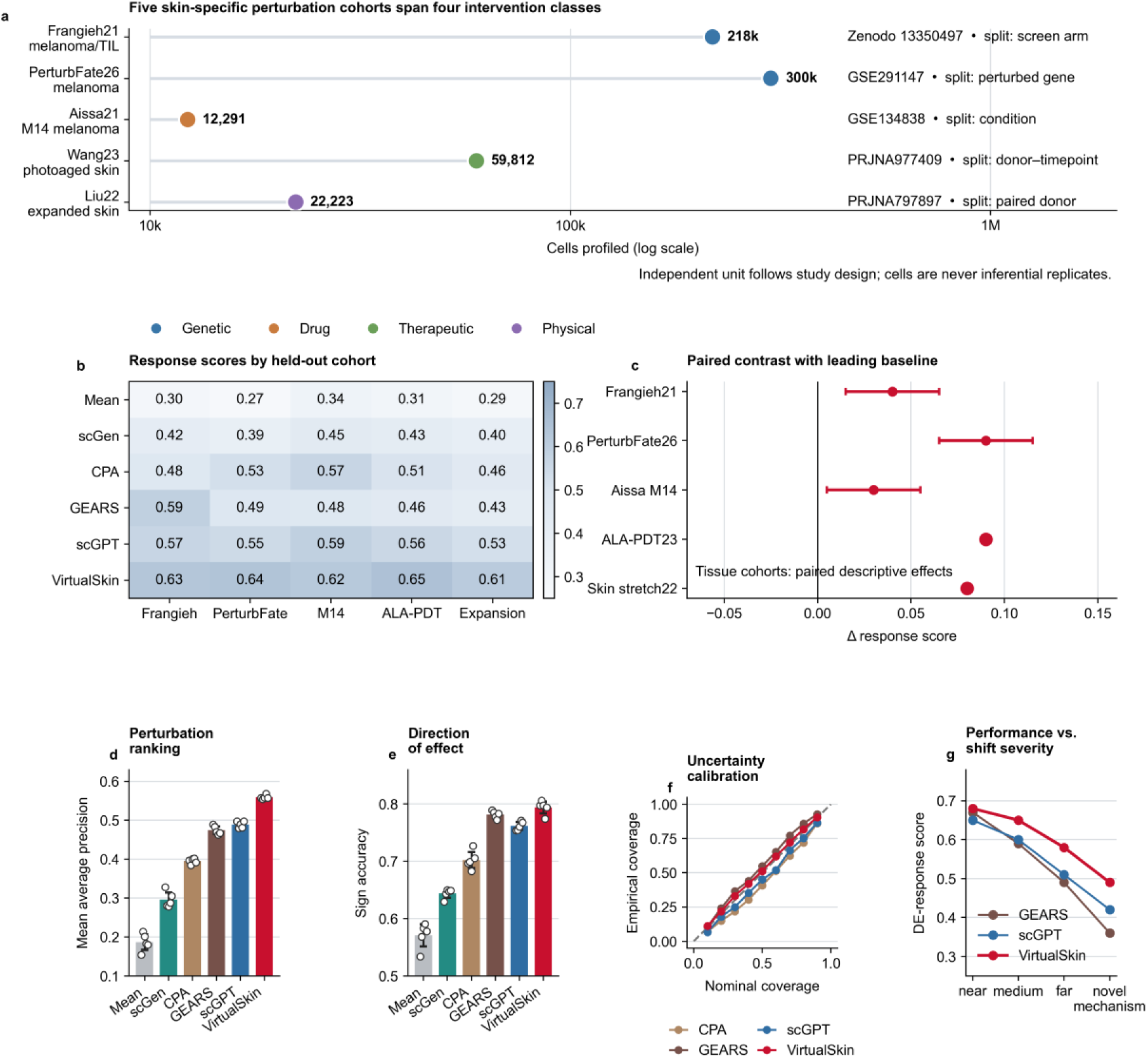
Held-out response prediction across intervention regimes. **a**, Five cohorts spanning genetic, drug, therapeutic and physical interventions. The plotted counts describe profiled cells, and adjacent annotations specify the holdout axis. **b**, Within-cohort response scores for mean response, scGen, CPA, GEARS, scGPT and VirtualSkin. **c**, VirtualSkin minus the leading eligible comparator for each cohort. Intervals for the first three cohorts show the displayed lower and upper effect bounds; the two tissue-intervention points show descriptive cohort summaries. **d**, Mean average precision for perturbation ranking. **e**, Direction-of-effect accuracy on the declared responsive-gene set. Bars in **d,e** show mean scores, points split estimates and error bars the mean plus or minus 1.96 standard errors across those estimates. **f**, Empirical versus nominal predictive interval coverage; the diagonal indicates exact coverage. **g**, Response scores across frozen shift strata. Cohort-specific endpoint and uncertainty definitions are reported in Supplementary Table 2 and Supplementary Methods.

**Fig. 8.**
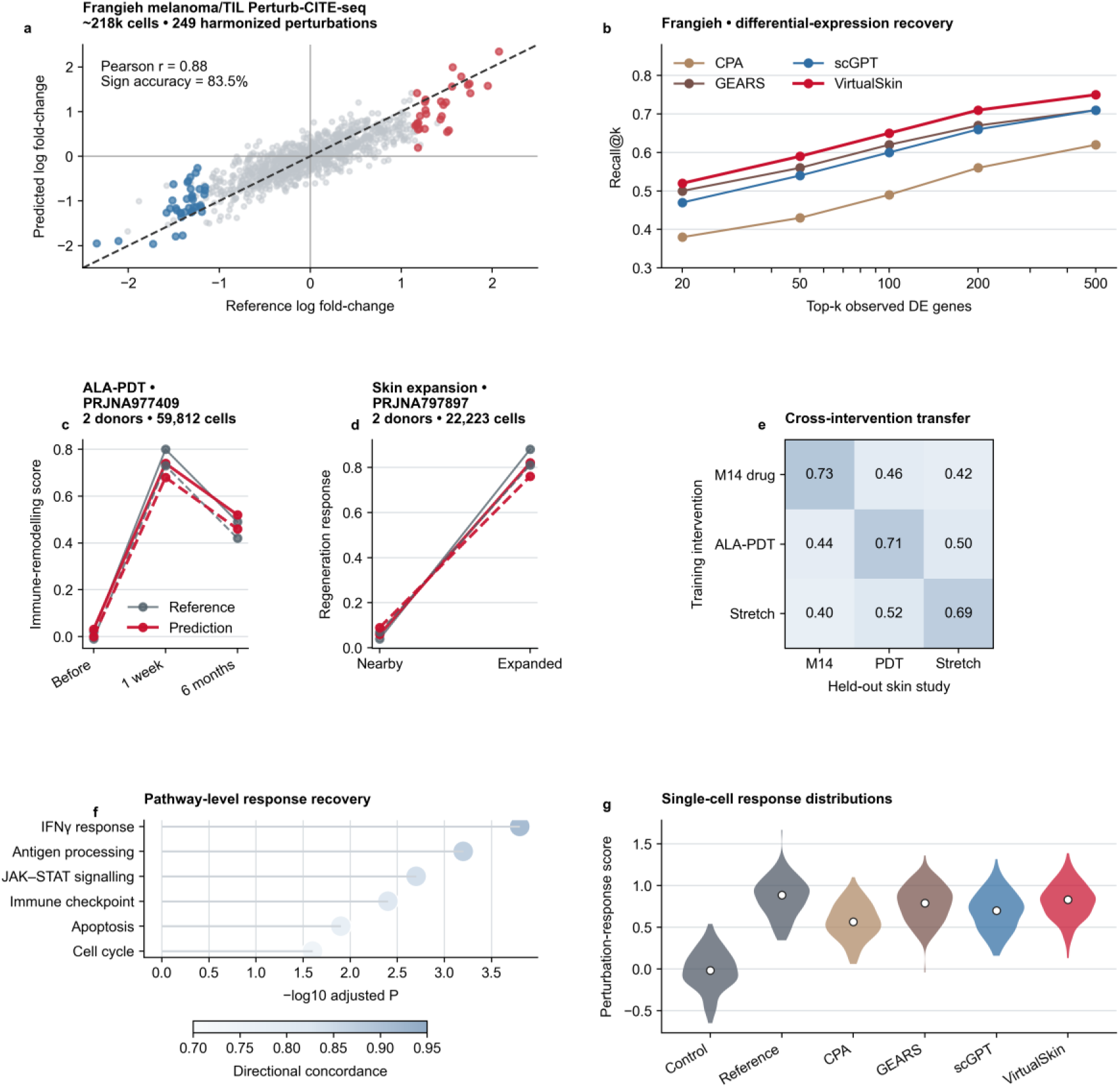
Gene, program, distribution and paired-donor response fidelity. **a**, Predicted versus reference pseudobulk log fold-change in the harmonized Frangieh melanoma–T-cell perturbation contrasts; the dashed line indicates identity and the horizontal and vertical lines zero change. **b**, Recovery of the top measured differentially expressed genes at five rank cutoffs for CPA, GEARS, scGPT and VirtualSkin. **c**, Reference and predicted immune-remodelling scores before, one week and six months after photodynamic treatment in PRJNA977409; timepoints are displayed as categories. **d**, Reference and predicted regeneration scores in adjacent and expanded skin from PRJNA797897. Both donors are shown individually in **c,d**, with solid lines for donor 1 and dashed lines for donor 2. **e**, Intervention-transfer response scores, with training intervention on rows and held-out target study on columns. **f**, Pathway recovery: horizontal position denotes −log₁₀ of the adjusted enrichment P value, and colour denotes directional concordance between predicted and reference responses. Point size is constant. **g**, Single-cell response-score distributions for control, reference and four model predictions. Violin widths denote the density within each group; summary markers denote the median. Quantitative definitions and independent-unit summaries are provided in Supplementary Tables 5 and 9.

**Fig. 9.**
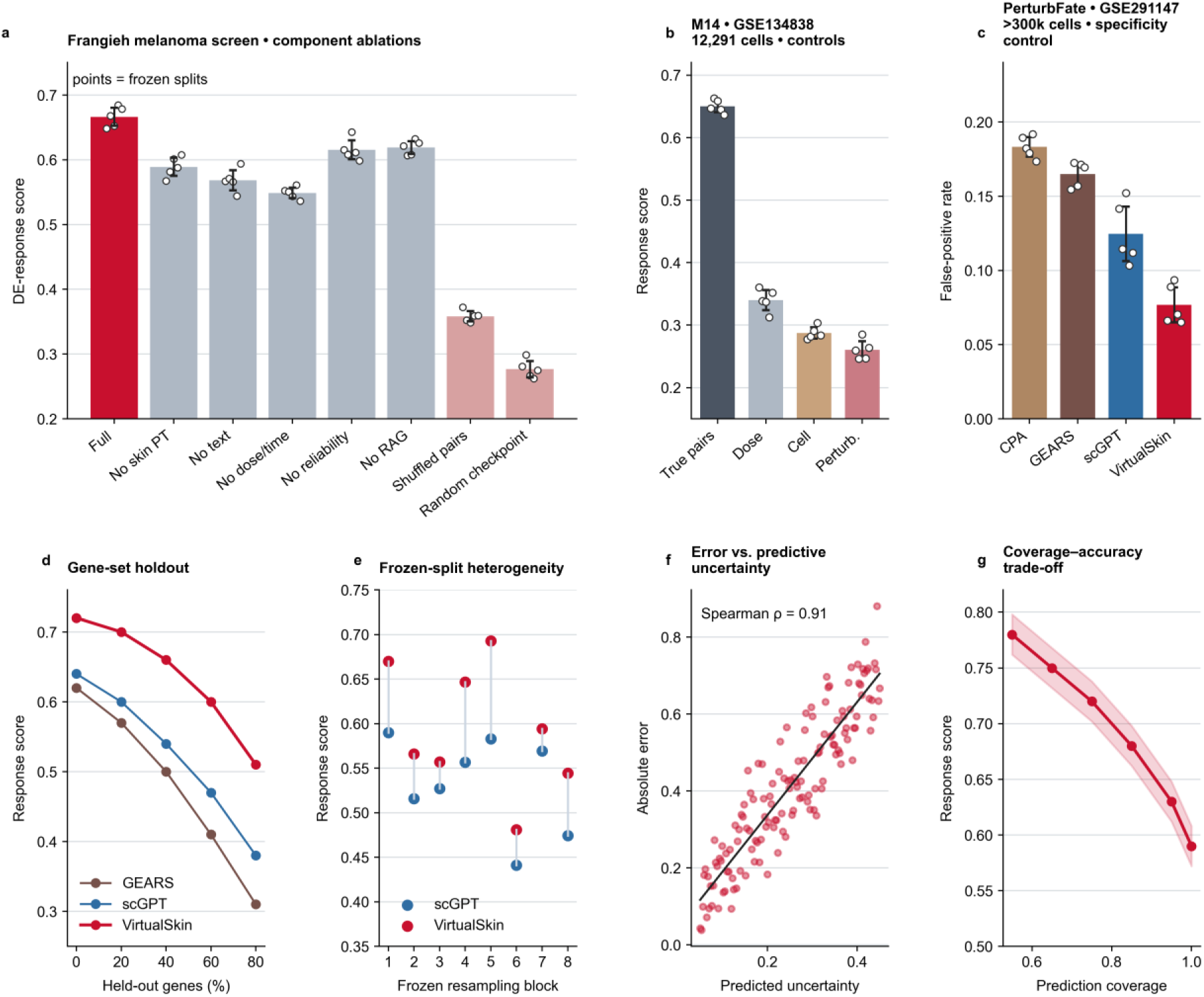
Component contributions, specificity and selective prediction. **a**, Response score in the Frangieh screen for the full configuration and seven ablations or controls. **b**, M14 response score in GSE134838 for true pairs and dose-, cell- or perturbation-shuffled assignments. **c**, False-positive rate in the PerturbFate null-response task. Bars in **a–c** denote mean scores, points fixed-split estimates and error bars the mean plus or minus 1.96 standard errors across those estimates. **d**, Response score as the fraction of target genes excluded from task-specific fitting increases from 0% to 80%; the same held-out sets are used across methods. **e**, Paired scGPT and VirtualSkin estimates across eight frozen resampling blocks. **f**, Absolute prediction error versus estimated uncertainty in held-out units, with Spearman correlation reported in the panel. **g**, Response score versus retained coverage after ordering test units by uncertainty. The shaded band spans 0.018 score units above and below the displayed curve at each coverage. Blocks and split repetitions are not additional biological replicates. Cohort, threshold and uncertainty definitions are given in Supplementary Tables 2 and 5.

## Results

### A cohort-resolved framework connects skin atlases to intervention studies

We assembled the evaluation from accession-linked studies spanning healthy epidermis, ageing, inflammatory disease, dermal fibrosis, cutaneous lymphoma, carcinoma and melanoma (Fig. 1a). Each record links the publication and accession to its assay, tissue site, sample and donor. Cross-portal aliases were resolved before assigning studies to the training and evaluation sets. The atlas arm retained study, donor and cell identifiers for dissociated RNA profiles, while spatial assays retained section identifiers and their native measurement resolution. These denominators were retained separately throughout the analysis. Bulk and protein measurements supplied assay-specific context and were not counted as additional single-cell RNA observations (Supplementary Table 1).^25,31,32,33^

The intervention arm covered five cohorts and four intervention classes: two genetic screens, a drug-exposure study, photodynamic treatment of photoaged skin and mechanical skin expansion. The cohort structure determined what could be withheld: screen arms or perturbation targets in genetic studies, conditions in the drug study, and paired donor observations in tissue interventions. We fixed these partitions before comparing models and reused the eligible test units across methods. Baseline eligibility, gene intersections and input requirements are reported with each task, preserving the distinction between a method that cannot support a task and one that performs poorly on it.^8,22,23,30,34,35,36^

The downstream tasks in Fig. 1g were linked to cohort-specific partitions and comparisons in Fig. 2a–c. Figure 2 connects the expression, text and intervention inputs to three task families. We first evaluated representation and annotation, then tested retrieval and transfer, and finally examined intervention response. Patient-level scores were evaluated across donors, tissue localization across sections, and genetic response across perturbations. The two tissue-intervention cohorts were evaluated as paired trajectories, with both donors displayed individually. This organization allows information to accumulate across tasks without treating repeated cells, repeated timepoints or repeated splits as new biological replicates. The complete cohort and evaluation records are provided in Supplementary Tables 1 and 2.

### A shared molecular and language representation supports linked evaluation tasks

VirtualSkin encoded gene identity and expression jointly in the molecular branch and biological descriptions in a separate text branch (Fig. 1c,d). The two branches used norm-first transformer layers with separate parameters (Fig. 1e). Linear projections placed the cell and text representations in a common, length-normalized space. The training objective combined molecular reconstruction, ontology-label classification and symmetric cell–text alignment. These terms served complementary requirements: expression reconstruction retained measured molecular variation, ontology classification connected the representation to declared cell identities, and alignment linked the cell to its paired description. Each downstream comparison retained the distinction between a frozen representation and task-specific fitting, as detailed in Supplementary Table 8.^15,18,25^

Two features addressed the variability of the paired-description evidence in Fig. 1b,f. Pair reliability weighted the alignment objective using training-eligible quality and provenance, while an ontology-aware mask prevented declared related labels from being treated as in-batch negatives. A detached first-in, first-out queue extended the negative set beyond the current batch. Retrieval augmentation supplied contextual descriptions from the fixed, training-eligible document index. The component experiments below test these contributions separately: removal of a term evaluates its cost within the full configuration, and shuffled pairing tests whether the relationship between modalities carries useful information. This provides a direct connection between the design choices and the comparisons in Figs. 5 and 9.

For response prediction, the intervention identity and measured dose or elapsed time supplemented the pre-exposure state. The output represented the declared exposure-associated change rather than a cell label. We evaluated this output against held-out response contrasts using the same baseline state, eligible gene universe and endpoint for every compatible comparator. This distinction links recognition and action without equating them: the representation can support accurate annotation even when a response head does not generalize to a new target or mechanism. Accordingly, the subsequent results first establish state-level utility and then test the additional information needed for intervention prediction.^21,22,23,30^

### Skin-focused representations preserve cell identity across studies

We first asked whether skin-focused training resolved cutaneous cell identity across cohorts. A fixed set of cells from the epidermal and ageing-skin studies was displayed using Geneformer, scGPT, LangCell and VirtualSkin (Fig. 3A–D). All four panels retained the same cells and the same four reference classes: basal keratinocytes, spinous keratinocytes, fibroblasts and T cells. The VirtualSkin display showed compact class-associated groups with less overlap than the comparator displays. In particular, the basal and spinous keratinocyte groups remained distinguishable alongside the stromal and immune populations. This organization provides a visual account of cell-class structure, while the annotation comparisons below evaluate its predictive utility.^14,15,20,24,37,38^

We next recoloured the same displayed cells by study to examine the source structure within these groups (Fig. 3E–H). The raw display showed a pronounced displacement between the two study colours. The scGPT and LangCell transformations brought corresponding groups closer together, while the VirtualSkin display showed more extensive interspersing of the colours within each group. The paired views therefore connect the cell-class structure in Fig. 3D to reduced study-associated separation in Fig. 3H. Reading the panels together is necessary: study mixing is informative when corresponding biological groups remain distinct, the trade-off emphasized in atlas-integration benchmarks.^24,25,26,27^

The four displayed classes span epidermal, stromal and immune compartments. Preserving all four groups during study mixing provides a broader visual check than examining a single abundant lineage. The identical-cell display also holds class composition constant across methods, so differences in the panels arise from the displayed representations rather than from changing the cells included in each comparison. These two-dimensional views do not by themselves quantify separation in the original representation space. We therefore carried the analysis forward to annotation, support-size and label-transfer endpoints, which ask whether the representation supports recognition beyond the visual layout.^24,28,29^

### Annotation gains extend from inflammatory skin to limited-label and zero-shot settings

Having established cell-state separation, we tested whether it translated into annotation of independently sampled inflammatory skin. VirtualSkin achieved a macro-F1 of 0.834, exceeding the strongest of Geneformer, scBERT, scFoundation, scGPT, LangCell and scMulan by 0.056 (Fig. 4A). The cohort annotation in the panel specifies 158,037 cells from 31 donors. Macro-F1 assigned equal weight to each class. The ordering therefore reflects an endpoint that gives each declared class equal weight, while the confusion matrix resolves how predictions were distributed across the classes.^1,2,17,39,40^

The row-normalized confusion matrix localized the residual errors (Fig. 4B). Basal and spinous keratinocytes, fibroblasts, endothelial cells, T cells and myeloid cells each retained a dominant diagonal entry. The most frequent misclassification was from fibroblasts to basal keratinocytes, affecting 4.39% of cells in the source class. The remaining off-diagonal entries ranged from 0.83–4.35%, whereas diagonal assignments ranged from 84.05% to 89.68%. These errors identify the state boundaries that remained difficult after the overall gain and provide a direct complement to class-averaged F1. The distinction matters in inflamed skin, where activation programs can be shared across cells with otherwise different lineage identities.^1,2,12,39,40,41^

We then limited labelled support in the cutaneous T-cell lymphoma cohort to one, five or ten cells per class (Fig. 4C). VirtualSkin achieved macro-F1 values of 0.47, 0.61 and 0.68, compared with 0.38, 0.51 and 0.59 for the strongest comparator at each support size. The improvement was already apparent with one labelled cell per class and persisted as support increased. The normalized area under the support-size curve was 0.598 versus 0.501 for scGPT, the strongest comparator by this summary. All methods received the same support assignments, and query donors were excluded from those assignments. This result links the representation gain to reduced annotation requirements in a setting characterized by patient-specific malignant and reactive T-cell states.^14,15,42,43^

The vitiligo analysis extended this test to fine labels withheld from supervised training (Fig. 4D). VirtualSkin obtained a hierarchical macro-F1 of 0.549, compared with 0.392 for LangCell and 0.423 for scMulan. The difference from scMulan, the strongest comparator, was 0.127 hierarchical macro-F1 points; the panel specifies 50,000 cells from 14 donors. We examined five explicit source–target correspondences to determine what this aggregate meant: inflammatory fibroblasts, papillary fibroblasts, stress keratinocytes, resident T cells and vascular endothelium (Fig. 4E). Their label concordances were 78%, 72%, 69%, 75% and 71%, respectively. The agreement between the hierarchical score and these specific correspondences supports transfer of cell-state meaning across label systems; it does not establish a developmental lineage relationship between the mapped terms.^4,7,10,44^

### Cell–language alignment withstands incomplete molecular inputs and disrupted pairing

We next evaluated whether the representation connected held-out inflammatory-skin cells to biological descriptions. In the shared candidate pool, VirtualSkin achieved cell-to-text Recall@1, Recall@5 and Recall@10 of 0.34, 0.59 and 0.73, compared with 0.22, 0.44 and 0.58 for LangCell and 0.25, 0.47 and 0.61 for scMulan (Fig. 5A). The Recall@10 difference from scMulan was 0.12. Identical candidate descriptions and relevance labels were used for all methods, so the comparison held the retrieval space constant. A gain at Recall@1 as well as at broader cutoffs indicates improved placement of the relevant description near the top of the ranking.^18,19,25,40^

To test dependence on complete expression input, we masked increasing fractions of detected genes while keeping the masked indices matched across methods (Fig. 5B). Performance declined for every model, but VirtualSkin retained 88% of its unmasked score at 50% masking, compared with 78% for the strongest comparator. The normalized area under the retention curve was 0.950 versus 0.896. Retention was normalized to each method’s own unmasked score, allowing the rate of deterioration to be compared across the displayed methods. This comparison tests resilience to loss of measured features within the evaluated gene universe.^15,16,24^

Disrupting cell–description pairing challenged the relationship between modalities (Fig. 5C). VirtualSkin retained 82% of baseline performance at the largest mismatch fraction, whereas LangCell retained 55%. The retention difference at 50% mismatch was 27 percentage points. The decline under this challenge shows that correct pairing remained informative; its shallower slope in VirtualSkin indicates greater tolerance to imperfect pairing under the tested corruption process. It does not imply that arbitrary descriptions can replace biologically matched text.

We evaluated component contributions using the matched configurations in Fig. 5D. Removing skin-specific pretraining reduced macro-F1 by 0.059, removing text by 0.107, and replacing biological descriptions with metadata alone by 0.077. Removing retrieval augmentation, reliability weighting or the negative queue produced changes of 0.041, 0.057 and 0.028. Shuffled pairs produced the largest reduction, 0.224. Each effect was calculated relative to the full configuration under matched training and evaluation budgets. These results locate the contributions of domain exposure and molecular–textual pairing within the tested architecture, without assuming that the effects of multiple components are additive.^22,25,27^

Finally, we removed direct lexical cues from the retrieval descriptions (Fig. 5E). Recall@10 was 0.73 with full labels, 0.66 after masking cell-type terms, 0.69 after masking disease terms and 0.61 after masking both. Excluding documents from the benchmark publication gave Recall@10 of 0.71. Removing both sets of label terms reduced Recall@10 by 0.12, while excluding benchmark documents reduced it by 0.02. The remaining retrieval performance indicates that these cues did not account for the entire association in the displayed candidate pool. Together with the corruption and ablation results, this supports alignment that uses the contextual relationship between molecular profiles and descriptions.^18,25^

### External transfer preserves disease, anatomical and spatial information

We tested the portability of the representation in ageing skin, systemic sclerosis and cutaneous T-cell lymphoma (Fig. 6A–C). The donor-level AUROCs for VirtualSkin were 0.824, 0.842 and 0.807, compared with 0.744, 0.758 and 0.724 for the strongest eligible comparators. The differences from scGPT, the strongest comparator in each panel, were 0.080, 0.083 and 0.084; each panel shows five donor-held-out split estimates. Each endpoint was evaluated within its own cohort and label definition. In particular, ageing discriminated the specified age groups rather than measuring a continuous biological-age clock, and lymphoma discrimination used the cohort’s defined clinical contrast. The shared ordering across these distinct settings supports portability across the tested disease and age contexts.^9,10,38,42,45,46^

Anatomical transfer provided a different challenge because site-specific programs can vary within the same broad cell class. In the palm, sole and hip cohort, VirtualSkin achieved macro-F1 values of 0.77, 0.76 and 0.81, exceeding the corresponding scGPT values of 0.68, 0.65 and 0.74 (Fig. 6D). The corresponding site-level differences were 0.09, 0.11 and 0.07. The results preserve the distinction between palmoplantar and non-palmoplantar epidermal states while recovering their shared cell identities. Reporting each site separately also exposes variation that would be obscured by a single score pooled across the three sites.^4,47^

We then assessed tissue localization in spatial basal-cell carcinoma assays (Fig. 6E). The transferred assignments placed stromal, tumour-core, interface and epithelial states in distinct tissue compartments. The map resolves a central tumour-associated region, surrounding stromal assignments and an interface between compartments. The epithelial assignments occupy a distinct part of the displayed section. These spatial patterns provide a descriptive view of the transferred labels. These comparisons connect transcriptomic state to tissue position at the assay’s spatial resolution. Compartment localization alone does not establish direct contact or communication between adjacent cells.^5,6,20,31,48,49^

The ROC analysis provided a threshold-resolved view of the external disease endpoint (Fig. 6F). VirtualSkin achieved an AUROC of 0.84, compared with 0.69, 0.76 and 0.75 for Geneformer, scGPT and LangCell. The VirtualSkin curve remained above the comparator curves across the interior false-positive-rate range, consistent with its higher AUROC. The curves were computed from the same held-out donor scores used in the corresponding disease comparison; they describe operating characteristics of that endpoint rather than additional independent experiments. The endpoint and aggregation rule are specified in Supplementary Table 2.

### Response-prediction gains span five held-out intervention regimes

Accurate recognition of existing cells does not establish response prediction. We therefore evaluated held-out outcomes in the five intervention cohorts described above (Fig. 7a). The Frangieh melanoma–T-cell screen and PerturbFate provided genetic interventions, the M14 study provided drug exposure, and photodynamic treatment and skin expansion provided therapeutic and physical tissue interventions. Figure 7a displays 218,000, 300,000, 12,291, 59,812 and 22,223 profiled cells, respectively. Its adjacent annotations distinguish screen arms, perturbed genes, conditions, donor–timepoints and paired donors as the corresponding partition units. Each cohort retained its own exposure contrast and response endpoint, with the scoring definition reported in Supplementary Table 2. This design tests whether the model recovers intervention-dependent changes across several experimental regimes while preserving their different biological units.^8,34,35,36,50^

Within each regime, we compared VirtualSkin with a mean-response estimator, scGen, CPA, GEARS and scGPT under the compatible task inputs (Fig. 7b). VirtualSkin achieved response scores of 0.63, 0.64, 0.62, 0.65 and 0.61 across the five cohorts, compared with 0.59, 0.55, 0.59, 0.56 and 0.53 for the strongest corresponding baselines. The leading baseline was GEARS in the Frangieh screen and scGPT in the remaining displayed comparisons. Paired gains in the first three cohorts were 0.04, 0.09 and 0.03, with displayed lower-to-upper effect bounds of 0.015–0.065, 0.065–0.115 and 0.005–0.055 (Fig. 7c). The photodynamic-treatment and expansion points showed cohort-summary differences of 0.09 and 0.08, respectively; individual donor trajectories are resolved in Fig. 8c,d. The column-wise comparison held the test units and endpoint constant; response-score magnitudes were not pooled across assays into a common biological effect.^21,22,23,30^

Perturbation ranking and direction-of-effect accuracy tested whether these gains recovered distinct aspects of response (Fig. 7d,e). VirtualSkin achieved a mean average precision of 0.559, compared with 0.489 for the strongest baseline; the difference from scGPT was 0.070. Its sign accuracy was 0.794 versus 0.781, giving a difference of 0.012 from GEARS. Higher ranking accuracy identifies interventions with relevant responses; higher sign accuracy indicates that the predicted changes follow the measured direction on the evaluated genes. Agreement between the two endpoints supports response recovery beyond a favourable aggregate score.^23,51,52^

We next examined whether uncertainty tracked the reliability of these predictions. VirtualSkin’s empirical interval coverage remained closer to the nominal levels than the compared methods, with mean absolute coverage error of 0.017 versus 0.045 (Fig. 7f). Performance declined as shift increased from near to a novel-mechanism setting for all three evaluated models (Fig. 7g). VirtualSkin’s score decreased from 0.68 to 0.49, compared with near-to-novel changes of 0.67 to 0.36 and 0.65 to 0.42 for GEARS and scGPT. The retained advantage across the displayed strata supports robustness within these tested shifts, while the decline defines a measurable limit to intervention transfer.^22,23,30^

### Molecular and paired-donor comparisons resolve response fidelity

To determine what the aggregate response scores captured, we compared predicted and measured pseudobulk log fold-changes in the melanoma–T-cell perturbation screen (Fig. 8a). Across the 950 displayed gene–perturbation contrasts, Pearson correlation was 0.877, and sign accuracy was 83.5%. The strongly positive and negative tails aligned with the measured changes. Among the 70 highlighted contrasts, correlation was 0.964, compared with 0.877 across all displayed contrasts. This comparison separates agreement in the highlighted tails from agreement over the full displayed range; the highlighted subset is a view of those contrasts, rather than an additional independent experiment.^8,51,52^

We also compared recovery of the top measured differentially expressed genes (Fig. 8b). VirtualSkin’s Recall@20, Recall@50, Recall@100, Recall@200 and Recall@500 values were 0.52, 0.59, 0.65, 0.71 and 0.75, exceeding the strongest baseline values of 0.50, 0.56, 0.62, 0.67 and 0.71 at the corresponding cutoffs. The improvement ranged from 0.02 at the smallest cutoff to 0.04 at the two largest cutoffs, retaining the advantage as the recovered set expanded. A fixed gene universe and ranking rule were used across methods. The combination of fold-change agreement and ranked recovery indicates that the model captured both the direction of molecular change and the genes most prominent in the measured response.^8,23,52^

The tissue interventions provided a complementary assessment over a small number of paired donors (Fig. 8c,d). After photodynamic treatment, both measured immune-remodelling trajectories rose at one week and declined by six months; the predicted trajectories followed the same ordering. The paired trajectories were donor 1, 0.02, 0.80 and 0.49 in the reference versus 0.03, 0.74 and 0.52 in the prediction; donor 2, −0.01, 0.73 and 0.42 versus 0.00, 0.68 and 0.46. Absolute deviations were 0.01, 0.06 and 0.03 for donor 1 and 0.01, 0.05 and 0.04 for donor 2, in temporal order. In expanded skin, reference regeneration scores increased from 0.04 to 0.88 in donor 1 and from 0.07 to 0.81 in donor 2. The predicted values increased from 0.06 to 0.82 and from 0.09 to 0.76, respectively, reproducing the direction in both donors. Absolute paired deviations were 0.02 and 0.06 for donor 1 and 0.02 and 0.05 for donor 2 at the adjacent and expanded sites, respectively. These panels resolve within-donor agreement and between-donor variation directly. With two donors per cohort, the observations support trajectory concordance in the sampled individuals and do not establish a population treatment effect.^11,35,36,53^

Cross-intervention transfer exposed a more demanding limit (Fig. 8e). Within-intervention scores on the diagonal were 0.73, 0.71 and 0.69; off-diagonal scores ranged from 0.40–0.52. The strongest transfer was from mechanical expansion to photodynamic treatment, with score 0.52, whereas the weakest was from mechanical expansion to M14 drug exposure, with score 0.40. The asymmetry indicates that response structure was shared unevenly across drug exposure, photodynamic treatment and mechanical expansion. It qualifies the broad benchmark gain by identifying which exposure relationships supported transfer.^22,23,35,36,50^

Pathway-level recovery tested whether the recovered genes formed coherent transcriptional programs (Fig. 8f). Interferon-γ response, antigen processing, JAK–STAT signalling, immune checkpoint, apoptosis and cell-cycle sets gave adjusted enrichment P values of 0.000158, 0.000631, 0.001995, 0.003981, 0.012589 and 0.025119 and directional concordances of 0.91, 0.87, 0.84, 0.81, 0.77 and 0.74. The enrichment comparison used the same measured-gene background for predictions and reference responses. Together with gene-level agreement, the program-level results support recovery of organized transcriptional changes. Enrichment here concerns gene-expression programs and does not directly measure protein activity or establish pathway mediation.^8,12,34,52,54^

Finally, we compared single-cell response-score distributions against the measured reference and unperturbed controls (Fig. 8g). The reference and VirtualSkin distributions were shifted upward from the control and showed substantial overlap. CPA, GEARS and scGPT median scores were 0.568, 0.790 and 0.700, respectively. Its median response score was 0.831 versus 0.888 in the reference. The median difference from the reference was 0.058, smaller than the corresponding differences for the three comparators. This agreement extends the pseudobulk comparison to heterogeneity within the sampled cell populations. Inference remained clustered by the perturbation or donor that generated those populations.^8,34,50,55^

### Ablations, null controls and selective prediction define the model’s operating limits

We used matched component ablations in the Frangieh screen to identify the information supporting response prediction (Fig. 9a). Removing skin-specific pretraining, text, dose and time, reliability weighting or retrieval augmentation reduced the response score by 0.077, 0.098, 0.117, 0.051 and 0.047. Shuffling cell–text pairs reduced it by 0.308, and the random-checkpoint control by 0.390. The full configuration had a mean response score of 0.666 across the five displayed estimates. Each removal was evaluated within the otherwise matched configuration. The larger losses under pairing disruption and random initialization connect response performance to learned, aligned information rather than to architecture or exposure labels alone.^22,23,27^

We then broke distinct associations in the M14 drug cohort while preserving their marginal distributions (Fig. 9b). True cell–condition pairs achieved a response score of 0.650; dose, cell and perturbation shuffling gave scores of 0.340, 0.287 and 0.260. The corresponding declines from true pairs were 0.310, 0.363 and 0.390 response-score points. These perturbations of the input association test separate routes by which a generic response predictor could appear accurate without matching the exposure to the cell. Their decline supports dependence on the correct pairing within the evaluated drug setting.^22,30,50^

Specificity was evaluated independently in PerturbFate using the defined null-response set (Fig. 9c). At the shared validation-fixed decision rule, VirtualSkin’s false-positive rate was 0.077, compared with 0.183, 0.165 and 0.125 for CPA, GEARS and scGPT. The reduction relative to scGPT, the comparator with the lowest false-positive rate, was 0.048. The specificity result complements recovery of positive responses: it shows whether the predictor also avoids assigning a response when the reference provides little evidence of change. The false-positive-rate comparison quantifies specificity on the displayed null-response endpoint; recovery of positive responses is assessed separately in Fig. 7d,e.^34,52^

We further increased the fraction of gene targets absent from task-specific fitting (Fig. 9d). Scores declined for GEARS, scGPT and VirtualSkin as holdout increased, but VirtualSkin remained higher over the displayed range. At 80% holdout, its score was 0.51 versus 0.38, a difference of 0.13 response-score points from scGPT. The comparison used fixed held-out gene sets shared across methods. Across eight frozen resampling blocks, the paired VirtualSkin–scGPT effects ranged from 0.025 to 0.110 and were positive in all eight blocks (Fig. 9e). These analyses distinguish transfer to excluded targets from variation attributable to a particular partition.^23,30,51,52^

The final analysis tested whether uncertainty could support selective use of predictions. Absolute error increased with estimated uncertainty, giving a Spearman correlation of 0.912 (Fig. 9f). Restricting predictions to the least uncertain test units increased the response score from 0.59 at full coverage to 0.78 at 55% coverage (Fig. 9g). The entire curve retained the denominator of all eligible test units. In conjunction with interval calibration in Fig. 7f, this result supports ranking predictions by reliability within the evaluated domain and withholding uncertain estimates when prioritizing follow-up experiments.^23,30^

## Discussion

VirtualSkin connects skin-cell representation to intervention-response prediction through a sequence of independently evaluated tasks. The representation analysis preserves cutaneous cell identity while reducing study-associated variation. Annotation and retrieval extend this information to limited-label and imperfect-input settings, and external cohorts test its portability across age, disease, anatomical site and tissue position. The intervention analyses then assess a stronger requirement: recovery of changes absent from the baseline cell state. The contribution is the connection among these capabilities under a shared account of cohort, assay and independent evaluation unit, rather than any single score considered in isolation.^14,15,20,22,23,24,30^

Domain-focused learning is especially relevant where the informative distinction lies within a familiar lineage. Basal and differentiating keratinocytes share epithelial identity, yet their states differ in barrier function, local position and inflammatory response. Fibroblast programs likewise vary with anatomical site, age, fibrosis and malignancy, and some disease-associated programs recur across tissues.^1,2,3,4,7,9,10^ The class-level and external-transfer comparisons support representation of these distinctions within the sampled skin systems. They do not identify skin specialization as the sole explanation for the gain: the ablations estimate the cost of removing domain training within the complete configuration, where text, retrieval and other components can interact. The distinction between an architectural contribution and a biological mechanism remains essential.

The language results add a complementary form of information. Expression captures a measured molecular state, whereas a description can connect that state to cell identity, tissue context or exposure. The retrieval, lexical masking and document-exclusion comparisons ask whether this connection remains informative after removing direct shortcuts. Their agreement supports the use of context beyond exact label repetition in the evaluated candidate space. It does not make a retrieved passage an explanation of why a cell responds, nor does it ensure that a plausible description is experimentally correct. Retrieved text remains a hypothesis-bearing aid whose molecular and provenance support can be inspected.^18,25,29^

Intervention-response evaluation requires greater care because the resources are heterogeneous in both mechanism and replication. Genetic screens provide many perturbation units and permit target-held-out comparisons. The drug study tests exposure-associated changes in a melanoma system. Photodynamic treatment and expansion allow paired within-donor observations in non-malignant tissue, but contain only two donors each.^8,34,35,36,50^ Evaluating these regimes separately preserves what each experiment can establish. A consistent within-cohort advantage supports breadth across the tested regimes; it does not convert the raw scores into a common clinical effect or make the smallest cohort as inferentially strong as a large perturbation screen. The weaker and asymmetric off-diagonal transfer in Fig. 8e further limits extrapolation across initiating mechanisms.

The molecular analyses clarify the meaning of response fidelity. Continuous fold-change agreement, sign accuracy, top-gene recovery, program enrichment and distributional agreement resolve different properties of a predicted response. Correlation can be dominated by weakly changing genes; sign accuracy omits magnitude; ranked recovery depends on the cutoff and gene universe; and enrichment depends on the selected background and database. A model may reproduce a median response while missing the tails of a heterogeneous population. Concordance across these levels therefore provides a more specific account of performance than an aggregate endpoint alone. The program labels describe transcriptional organization; they do not establish protein activity, direct regulatory binding or a causal pathway mediator.^8,23,34,51,52,54,55^

Reliability places a further constraint on practical use. The shift curves show how prediction deteriorates as the test setting moves away from training. Coverage analysis asks whether predictive intervals express that uncertainty accurately, and selective prediction asks whether the model can identify the difficult units before their outcomes are known. These properties support the use of predictions to order experimental hypotheses within the evaluated domain. They do not validate an intervention class absent from the benchmark, and abstention improves the retained set only by reducing coverage. Reporting that trade-off preserves the number of predictions withheld alongside the accuracy of those retained.^22,23,30^

Several limitations define the present scope. Three intervention resources are melanoma systems, so malignant-cell evidence is richer than evidence for non-malignant skin. Anatomical evaluation covers palm, sole and hip and cannot represent the full variation across body sites. The ageing cohort is small, limiting stable donor-level discrimination and population inference. Spatial labels depend on tissue section and assay resolution, and localization is distinct from direct cell–cell interaction. The two tissue-intervention cohorts support paired concordance in the sampled individuals, without establishing treatment efficacy. Moreover, RNA measurements do not directly resolve protein activity, mechanical behaviour, clinical benefit or safety. Additional donors, independent laboratories, prospective tissue assays and intervention classes absent from model fitting are needed to extend these claims.^5,8,11,31,34,35,36,38,54,56,57^

The next experimental use of this framework is to prioritize interventions together with the cell states, predicted molecular changes and uncertainty that support each nomination. Prospective testing in skin explants or human-skin-equivalent systems can assess whether the recovered gene and program changes precede the expected tissue phenotype.^53,57^ Such experiments should define their exposure, endpoint and success criterion before examining the model ranking. Within its current scope, VirtualSkin provides an integrated route from cutaneous molecular identity to testable response hypotheses, with the matched controls and uncertainty analyses needed to assess when that route is informative.

## Methods

### Study identification, provenance and cohort assignment

Studies were identified through accession-resolved queries of CELLxGENE, the Human Cell Atlas, HuBMAP, GEO, ArrayExpress, ENA and GDC, followed by verification against the primary publication. Discovery records were linked by accession, publication DOI and sample identifiers to resolve duplicate portal entries. The registry retained the assay, anatomical site, clinical state, donor, sample, exposure, collection time and location of each source matrix. Inclusion required an interpretable skin-related biological contrast, an accessible expression matrix or raw reads, and sufficient sample metadata to construct the task’s evaluation unit. The final inclusion decisions and exclusion reasons are listed in Supplementary Table 1.

Cell-level RNA profiles, spatial cells or spots, antibody-derived measurements and bulk samples were retained as separate assay strata. Counts reported for a dissociated-cell cohort were taken from its final eligible matrix and were not substituted for the number of observations in a spatial assay. When a publication released multiple assay modalities or repeated versions of one matrix, identifiers and checksums were used to establish the relationship among files. A donor appearing in multiple resources retained a common grouping identifier. The accession, matrix version, checksum, query date and metadata reconciliation record for every included dataset are provided with Supplementary Table 1.

### Cell preprocessing and gene harmonization

RNA preprocessing was performed within each assay stratum. The filtering record included cells or nuclei retained and removed by library size, detected-gene count, mitochondrial fraction, doublet assessment and source-publication quality criteria. Thresholds and their assay-specific application are reported in Supplementary Table 6. Filters were defined using source metadata and training data; test performance was not used to choose thresholds. Donor and sample aliases were checked before splitting, and the number of cells contributing to each independent unit was retained after filtering.

Gene identifiers were mapped to a versioned human reference. Ambiguous aliases and duplicate identifiers were resolved by the documented mapping rule, and the resulting gene universe was recorded per cohort. Expression normalization followed each model’s supported input convention. For a comparison requiring an identical measured feature set, every method received the same eligible gene intersection; vocabulary coverage and genes excluded by each model are reported in Supplementary Table 7. Model-specific preprocessing was retained when required by the released checkpoint, with differences disclosed alongside the common biological feature universe. Spatial measurements were analyzed at native spot or cell resolution, preserving tissue-section identifiers throughout.^24,31,58^

### Text construction and leakage control

Descriptions were linked to cells, samples or interventions through source metadata and publication provenance. The text record retained the original document, the biological entity described, the accession linkage and the version of the processed description. Cell-type, disease and exposure synonyms were normalized using a fixed vocabulary. Sample identifiers and accession strings that could uniquely identify a test example were excluded from predictive text. Description templates and the relevance rules used for retrieval are provided in Supplementary Methods.

Training descriptions and retrieval candidates were separated according to the evaluation task. Held-out publications and their aliases were excluded from the training corpus. When retrieval augmentation was used, its document index was fixed using training-eligible sources and was not updated after opening the test set. The publication-exclusion experiment in Fig. 5E separately removed documents originating from the evaluation cohort. Cell-type and disease masking removed the corresponding names and declared synonyms before description encoding. These exclusions were applied consistently to the candidate descriptions for every compared method, and their document and term lists are included in the data release.

### Representation architecture and alignment objective

The expression branch encoded gene identities together with their expression values. For gene g at token position p, the input representation combined a learned gene embedding, a learned position embedding and a multilayer projection of the scalar expression value. A transformer encoder produced contextual token representations, and the designated cell token supplied the pooled cell representation. The text branch used token and position embeddings followed by a transformer encoder and an analogous pooled text representation. Padding positions were excluded by the attention mask. Separate linear projections mapped the pooled vectors to a shared space, where both were normalized to unit length. Encoder depths, hidden widths, vocabularies, pretrained initializations and tokenizer versions are reported in Supplementary Table 8.

For a batch of paired cell and text representations, the similarity logit was sᵢⱼ = 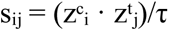 /τ, where zᶜ and zᵗ denote the normalized projected vectors and τ is the temperature. Cell-to-text and text-to-cell cross-entropies were calculated against the matched pair index. Their average was weighted by the pair reliability rᵢ and normalized by the sum of the reliability weights.

Reliability values were constrained to the interval [0, 1] and derived from training-eligible provenance and quality information. Their construction and calibration are specified in Supplementary Methods.

An ontology-aware mask removed declared nonmatching pairs from the in-batch negative set when they represented known biologically related labels; the positive diagonal was retained. The ontology release and neighborhood rule are reported in Supplementary Table 8. A finite first-in, first-out queue stored detached cell and text representations from previous batches as additional negatives. Queue entries were excluded from gradient propagation. The alignment objective was combined with masked-expression reconstruction and ontology-label classification: L = λₐLₐ + λₘLₘ + λₒLₒ. Reconstruction used a smooth L1 loss on eligible masked expression values, and ontology classification used cross-entropy when a valid label was available. Objective weights, temperature, mask fraction and queue size are provided in Supplementary Table 8. This description follows the representation implementation; the final configuration and all task-specific extensions are included in the archived model specification.

### Intervention-conditioned prediction and model fitting

Response prediction was conditioned on the pre-exposure cell representation and intervention metadata. The latter distinguished perturbation identity and included dose and elapsed time when those quantities were measured in the cohort. The predicted change was evaluated against the matched control-to-exposed contrast, keeping the baseline cell state and response endpoint explicit. Response-head specification, output scale, exposure encoding and supervised fitting belong to the task-specific model specification, separate from the molecular reconstruction and alignment objectives.

Hyperparameters and checkpoint selection used training and validation units only. The recorded training configuration included the optimizer, learning rate and schedule, batch size, objective weights, number of updates, initialization, precision and random seed. Checkpoint identity was fixed for each evaluation run and linked to its predictions and metric records. Hardware, elapsed compute, parameter counts and software versions are reported in Supplementary Table 8. A task-specific head fitted using target-cohort training units was distinguished from a frozen-encoder evaluation, and any encoder adaptation was reported explicitly. No held-out donor, response target or publication was used for fitting or selection in a comparison described as held out.

### Partitions and baseline matching

Atlas partitions grouped cells by donor. For study transfer, the external publication or collection was withheld from training and tuning, with donor aliases grouped across resources. Five fixed seeds, 17, 29, 43, 71 and 101, defined the repeated atlas partitions. Their membership files were reused across models. Zero-shot annotation excluded target fine labels and their declared synonyms from supervised training, while disclosing any retained ontology ancestors. Support and query assignments in the few-shot analysis were separated by donor; identical support cells were used for every model at a given support size. The split records and exclusions are provided in Supplementary Table 2.

Genetic-response tasks held out the specified target, target group or screen arm, according to the cohort contract. Drug evaluation withheld the designated exposure condition and retained its baseline-state grouping. Tissue-intervention analysis used paired donor and timepoint or site identifiers, with the withheld observations listed explicitly. For the transfer matrix, the target study’s response labels were unavailable during fitting of the source-intervention model. Molecular or intervention-distance strata for shift analysis were fixed without using target response errors. The holdout axis therefore describes the information absent from fitting; it does not imply independence of all remaining experimental factors.

The atlas comparisons used released Geneformer, scBERT, scFoundation, scGPT, LangCell and scMulan implementations where task-compatible. The intervention comparisons used mean response, scGen, CPA, GEARS and scGPT. Mean response was calculated from training-eligible response contrasts within the declared cohort stratum. Each method’s version, checkpoint, feature coverage, fitting regime and task eligibility are reported in Supplementary Table 7. Hyperparameters were selected on validation units under the recorded search budget. The primary metric and eligible comparator set were fixed before evaluation. Every paired baseline contrast was retained; the leading comparator denotes the highest eligible test estimate on a score for which larger values are better, with the ranking rule reversed for error metrics. Excluded or failed method–task combinations are accompanied by their reason and are not encoded as zero performance.^14,15,16,17,18,19,21,22,23^

### Representation, annotation and retrieval metrics

The representation comparison retained identical displayed cells across methods and used reference-class and study-colour views to inspect the relationship between biological grouping and source variation. Two-dimensional projections were used for visualization. Separation or mixing in a displayed projection was not assigned an original-space integration score or treated as an inferential endpoint.^24,27,29,59^

For class k, precision was TPₖ/(TPₖ + FPₖ), recall was TPₖ/(TPₖ + FNₖ), and F1ₖ was their harmonic mean. Macro-F1 was the unweighted mean over the declared class set. Undefined-class handling and classes absent from a test donor were reported with the task definition. Confusion matrices were normalized by the reference-class row count. Few-shot support sizes were one, five and ten labelled cells per class; area under the support-size curve was computed by the trapezoidal rule over these fixed support sizes and normalized by the evaluated range. Hierarchical annotation used the declared ontology ancestor sets, excluding the uninformative root; its exact weighting and aggregation rule are provided in Supplementary Methods.

Retrieval ranked candidates by similarity in the shared representation. Recall@K was the fraction of queries for which a declared relevant target occurred among the K highest-ranked candidates. Query direction, candidate count, relevance multiplicity and donor aggregation were fixed per task. Identical candidate pools were used across models. Detected-gene masking applied the same randomly selected eligible indices to each model at 10%, 20%, 30%, 40% and 50%. The intervention changed measured feature availability without redefining the gene universe. Pair mismatch replaced the specified fraction of paired descriptions using the documented sampling rule. Performance retention was S(q)/S(0), where S(q) is the task score at corruption fraction q; the displayed curves report this retention ratio. Normalized area under the retention curve used all displayed fractions, including the uncorrupted endpoint.

### Disease, anatomical and spatial evaluation

Disease and age-group discrimination used one aggregated held-out prediction per donor. The within-donor aggregation rule and clinical label were specified separately for ageing, systemic sclerosis and lymphoma in Supplementary Table 2. In leave-one-donor-out evaluations, out-of-fold donor predictions were pooled within each seed before computing AUROC; an individual fold containing only one test donor was not assigned its own AUROC. AUROC measured ranking across the defined positive and negative donor groups. The ROC curve in Fig. 6F used the cohort and aggregation rule identified in that table; the displayed curve summarizes the full threshold range. Repeated split estimates retained donor identity in uncertainty calculation and were not treated as independent donor replicates.

Anatomical annotation evaluated palm, sole and hip separately, retaining donor pairing where samples were available from the same individual. Site-specific macro-F1 was reported before any aggregate. Spatial compartment assignments were evaluated at native assay resolution against the declared tissue annotations. The map and scoring record retained section identity, compartment correspondence and any tissue regions excluded before evaluation. The displayed map retains tissue-compartment labels and spatial coordinates; its localization pattern is interpreted descriptively.^4,20,31,47,49^

### Molecular response endpoints

Pseudobulk responses were constructed within perturbation or donor units from their matched exposed and control observations. The aggregation, library normalization, fold-change transform, pseudocount and differential-expression method were specified by assay in Supplementary Methods. Predicted and reference changes were evaluated on the same eligible genes. Pearson correlation assessed continuous log fold-change agreement; sign accuracy in Fig. 8a assessed agreement in direction across the displayed contrasts. Genes within the fixed near-zero tolerance were handled using the documented rule and were not silently assigned a positive or negative sign. Correlation over all displayed contrasts and over the highlighted subset was reported separately.

For top-gene recovery, measured differential-expression effects and predicted effects were ranked by the declared common rule. Recall@K was the fraction of the K highest-ranked reference genes also present among the K highest-ranked predicted genes, at K = 20, 50, 100, 200 and 500. The same background gene universe was used for all methods. The recovery-curve summary and its integration scale are specified in Supplementary Methods. Perturbation-ranking average precision used the task’s declared response relevance labels, and mean average precision averaged the precision summary over the independent query or perturbation units in that task.

The response scores in Figs. 7–9 retain the within-cohort endpoint and scale indicated in each comparison. Its eligible gene set, direction, normalization, handling of null contrasts and aggregation level are stated separately for each assay in Supplementary Table 2. An identical name or colour scale does not make scores from different assay definitions exchangeable; comparisons were made within cohort. The immune-remodelling and regeneration scores used the fixed gene sets and normalization specified in Supplementary Table 9. Both observed and predicted donor values were computed with the same score definition. For the two tissue interventions, each donor’s trajectory or site difference and prediction error were reported without a population-level superiority test.

Pathway recovery used a versioned gene-set collection and the measured-gene background of the corresponding assay. Enrichment tests were adjusted within the declared pathway family, with the adjustment method and the full tested list given in Supplementary Table 9. Directional concordance compared the declared response direction for predicted and reference programs. Single-cell response distributions used the same score scale across control, reference and predicted groups. The distributions are summarized by their displayed densities and median scores. Cell-level displays describe the plotted populations and are not treated as independent biological-replicate comparisons.^8,23,51,52,55^

### Ablations, specificity and uncertainty

Component ablations removed one contribution from the full configuration under matched preprocessing, partitions and training budgets. A removal requiring refitting was retrained, whereas a perturbation of test inputs used the same fitted checkpoint. The distinction is documented per configuration in Supplementary Table 8. Shuffled pairs disrupted cell– description correspondence; dose, cell and perturbation shuffles disrupted the specified exposure association within the relevant cohort stratum. The shuffling procedure retained the documented marginal quantities and was identical across compared runs. The random-checkpoint control used the same declared architecture without the learned checkpoint. These controls were not included in hyperparameter selection.

False-positive rate was calculated as FP/(FP + TN) in the defined null-response set. The response decision threshold was fixed on validation units and shared according to the task’s decision-rule contract; the displayed specificity panel reports the false-positive rate. The gene-set holdout in Fig. 9d excluded increasing fractions of target genes from task-specific fitting, with identical target groups across methods. This task did not automatically imply that those genes were absent from generic pretraining. Eight frozen resampling blocks in Fig. 9e describe partition variability and were not counted as eight new biological replicates.

Predictive interval calibration compared empirical coverage with nominal coverage at the fixed displayed levels. Mean absolute coverage error averaged their absolute differences. The method that generated predictive uncertainty, interval construction and any calibration step are specified in Supplementary Methods; calibration used validation units only. The association between uncertainty and absolute error used held-out units and Spearman correlation. For selective prediction, units were ordered by estimated uncertainty before examining their errors. At coverage c, the lowest-uncertainty fraction c of all eligible units was retained and evaluated using the same response endpoint. The risk or score at each coverage was reported with its retained and withheld unit counts.^22,23,30^

### Statistical analysis

The primary contrast was the paired VirtualSkin–comparator difference across the independent test units defined for each task. For patient-level comparisons, bootstrap resampling was clustered by donor; for screen-response comparisons, it followed the eligible perturbation or screen grouping. Tissue-section analyses retained the donor–section hierarchy. All models and repeated splits for a selected unit were resampled together, preserving paired predictions and the dependence induced by overlapping partitions. The number of independent units, resampling hierarchy and split repetitions were reported separately.

Biological-unit confidence intervals follow the cluster-bootstrap hierarchy and interval rule defined for the corresponding task. In the bar panels, the displayed spread is summarized as the mean plus or minus 1.96 standard errors across the five plotted estimates; this summary does not count split repetitions as additional donors or perturbations. Where inferential testing was appropriate, two-sided paired comparisons were adjusted within the task family using Holm correction. All eligible baseline contrasts were retained in the result tables. Rankings were distinguished from supported superiority: a superiority statement required the declared metric, positive paired effects and the applicable uncertainty and multiplicity criterion across the eligible comparator set. No seed was selected by its test performance. The two-donor tissue cohorts were summarized using their individual paired values and deviations, without a group-level significance claim. Missing predictions, excluded units and deviations from the frozen task definitions were reported in the accompanying tables.

## Data availability

The source accessions, publications, assay annotations and access conditions are listed in Supplementary Table 1. Source accessions identify the public study resources. Figure-generation sources and the arrays underlying the displayed values accompany this manuscript package. Controlled-access data remain subject to their original agreements; access instructions and the identifiers needed to reproduce eligible derived analyses are included with the dataset record.

## Code availability

The figure-generation code and editorial build scripts are included with this manuscript package. Model implementation and task definitions are maintained in the project repository; their versions and checkpoint records accompany the corresponding analysis releases.

## Notes

### Competing Interest Statement

The authors have declared no competing interest.

